# Endogenous BAX and BAK form mosaic rings of variable size and composition on apoptotic mitochondria

**DOI:** 10.1101/2023.08.18.553869

**Authors:** Sarah Vanessa Schweighofer, Daniel C. Jans, Jan Keller-Findeisen, Anne Folmeg, Mark Bates, Stefan Jakobs

## Abstract

One hallmark of apoptosis is the oligomerization of BAX and BAK to form a pore in the mitochondrial outer membrane, which mediates the release of pro-apoptotic intermembrane space proteins into the cytosol. Cells overexpressing BAX or BAK fusion proteins are a powerful model system to study the dynamics and localization of these proteins in cells. However, it is unclear whether overexpressed BAX and BAK form the same ultrastructural assemblies following the same temporal hierarchy as endogenously expressed proteins. Combining live– and fixed-cell STED super-resolution microscopy, we show that overexpression of BAK results in novel BAK structures, which are virtually absent in non-overexpressing apoptotic cells. We further demonstrate that in wildtype cells, BAK is recruited to apoptotic pores before BAX. Both proteins together form unordered, mosaic rings on apoptotic mitochondria in immortalized cell culture models as well as in human primary cells. In BAX– or BAK-single-knockout cells, the remaining protein is able to form rings independently. The heterogeneous nature of these rings corroborates the toroidal apoptotic pore model.

## Introduction

Apoptosis, a form of programmed cell death, is essential for a multitude of processes including development and tissue homeostasis (1). Apoptosis dysregulation contributes to a variety of diseases, including cancer, autoimmune disorders, and neurodegenerative diseases (2–4). The intrinsic apoptosis pathway leads to the activation of the pro-apoptotic BCL-2 proteins BAX and BAK, resulting in mitochondrial outer membrane permeabilization (MOMP) (5, 6). The consequent release of proteins into the cytosol from the mitochondrial intermembrane space (IMS) such as cytochrome *C* or Smac/DIABLO triggers the activation of caspases and ultimately results in cell death (7, 8).

The two apoptosis effector proteins BAX and BAK are structurally similar and functionally redundant (9, 10). To impair apoptosis, both proteins must be knocked-out (KO) (11) and the majority of BAX/BAK-double-KO mice die perinatally (12). BAX and BAK are differentially expressed, with most tissues containing higher levels of BAX than BAK (13). At the cellular level, the two proteins differ primarily in their subcellular localization, because they are shuttled between the cytoplasm and the mitochondrial outer membrane (MOM) at different rates (14). As a result, in healthy cells, the majority of BAK is localized at the mitochondria, while the majority of BAX is localized in the cytosol. Upon activation during apoptosis, BAX translocates to the MOM and both BAX and BAK undergo conformational changes that result in the formation of the apoptotic pore (15). The precise nature of the apoptotic BAX/BAK pore as well as the sequence of integration of the proteins into the pore remains ill-defined, limiting the possibility of pharmaceutically targeting BAX and/or BAK in dysregulated apoptosis (16).

Apoptosis has long been considered to be immunologically silent and for the release of the mitochondrial IMS proteins only minimal BAX activation has been shown to be necessary (17). In fact, a pore consisting of only 6 BAX molecules and a size of 6 nm would be sufficient to release a cytochrome C molecule (18). Still, it has been shown that BAX and BAK form rings lining large apoptotic pores (19–21). These pores lead to mitochondrial inner membrane herniation and allow the release of mitochondrial DNA (mtDNA) into the cytoplasm (22, 23) triggering an inflammatory response if caspase activation is inhibited. In BAX/BAK DKO cells, overexpressed BAX and BAK exhibit different assembly kinetics and the interplay of the two proteins was shown to tune the kinetics of mtDNA release (21). Yet, little is known about the temporal dynamics of BAX/BAK pore growth on the nanoscale as well as about the composition of the pore at endogenous BAX/BAK expression levels.

Therefore, in this study, we combined live– and fixed-cell STED super-resolution microscopy to investigate the temporal dynamics and performed an in-depth analysis of the spatial arrangement of BAX and BAK in the apoptotic pore. Our results support the idea of a growing apoptotic pore delineated by heterogeneous rings composed of both BAX and BAK, with BAK integration on average preceding the integration of BAX.

## Results

### BAX and BAK pores exhibit different spatial and temporal dynamics

Although the oligomerization of BAX and BAK into higher-order structures leading to pore formation and MOMP has been intensely investigated (19–21, 24, 25), the nanoscopic details of BAX and BAK pore growth and dynamics *in cellulo* remain largely elusive due to the small size of these structures. To study the spatial and temporal dynamics of BAX and BAK, we performed long-term live-cell STED super-resolution microscopy in apoptotic cells.

We transiently expressed BAX fused N-terminally to a HaloTag in U-2 OS wildtype (WT) cells stably overexpressing the MOM marker Snap-OMP25 [Fig 1A]. Cells were imaged in the presence of the caspase inhibitor QVD-OPh (26) to prevent the detachment of apoptotic cells from the coverslip. Live-cell STED microscopy recordings showed that overexpression of Halo-BAX alone was sufficient for the cells to undergo apoptosis, as non-transfected cells in the same sample did not undergo apoptosis [Fig 1A, upper right corner]. BAX rings developing from distinct BAX clusters continuously increased in size on fragmented and rounded mitochondria [Fig 1A, arrows, and FigS1A], usually reaching a diameter limited by the diameter of the mitochondrial fragment they were located on within 3-15 minutes [FigS1B].

**Fig 1.**
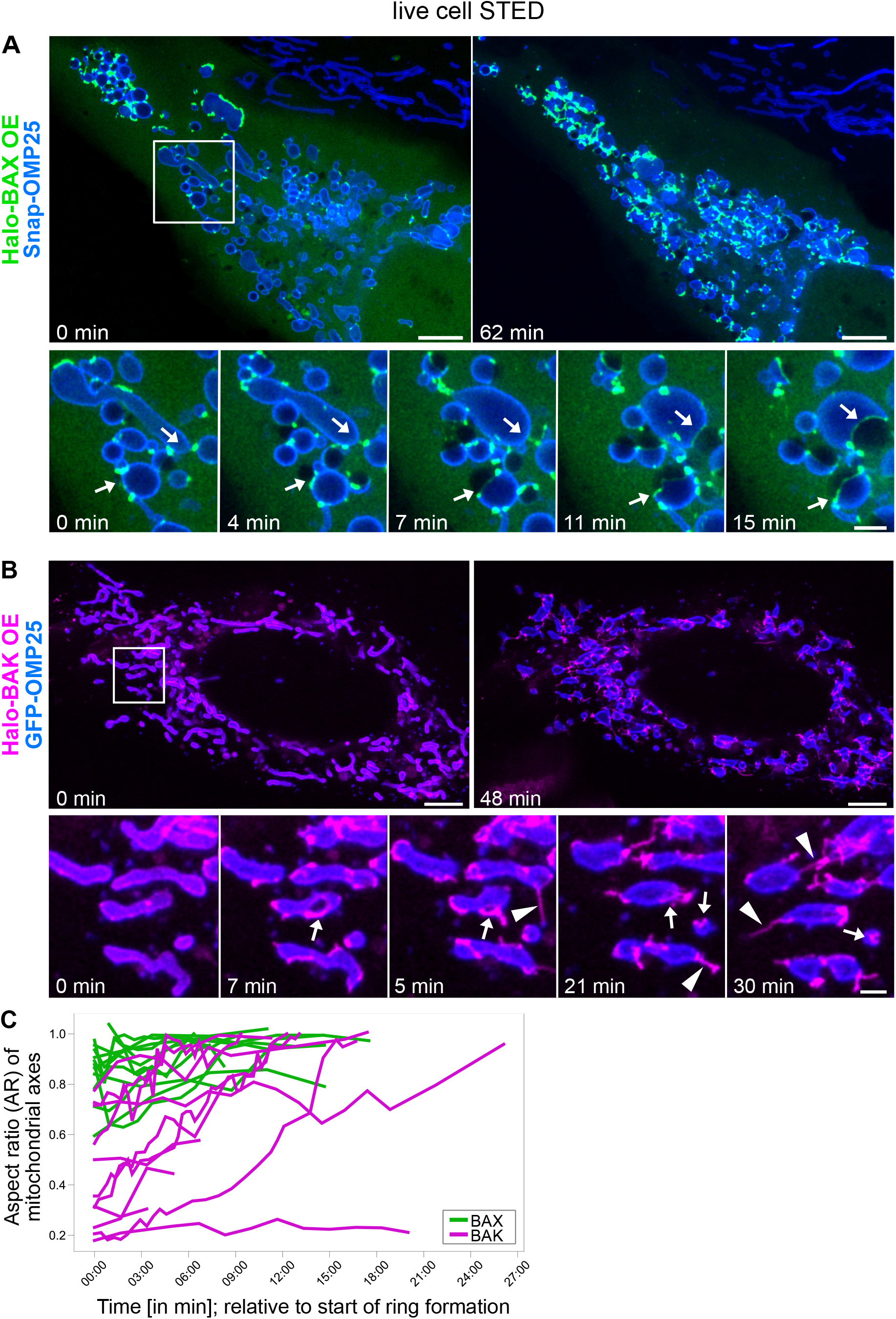
BAX and BAK pores display differences in spatial and temporal dynamics. (**A**) Upper panels: The first and last frames of a live-cell STED movie from a U-2 OS WT cell undergoing apoptosis. The cells stably overexpress Snap-OMP25 (labeled with silicone-rhodamine (SiR), blue) and were transiently transfected with Halo-BAX (labeled with Atto590, green). Cells were imaged in the presence of 20 µM QVD-OPh. Lower panels: Selected frames of the indicated area in the overview. BAX rings enlarge continuously on fragmented mitochondria (arrows). (**B**) Upper panels: The first and last frames of a live-cell STED movie from a U-2 OS WT cell undergoing apoptosis. The cells stably overexpress GFP-OMP25 (blue, confocal imaging mode) and were transiently transfected with Halo-BAK (labeled with Atto590, magenta). Cells were imaged in the presence of 20 µM QVD-OPh. Lower panels: Selected frames of the indicated area in the overview. BAK rings enlarge on fragmented but partially still tubular mitochondria (arrows). BAK “linkers” are devoid of MOM signal (arrowheads). (**C**) AR (^short mitochondrial axis^/_long mitochondrial axis_) of the mitochondrial fragments harboring BAX (green) or BAK (magenta) rings were plotted over time as a measure for mitochondrial shape dynamics. Data are quantified from 23 individual rings in 3 biological replicates of 2 independent experiments per condition. Scale bars: 5 µm (A and B, upper panels), 1 µm (A and B, lower panels).

Similarly, the overexpression of Halo-BAK [Fig 1B] was sufficient to induce apoptosis. BAK formed rings [Fig 1B, arrows], most of which enlarged continuously for up to 20 min, but remained smaller than the BAX rings [FigS1A]. Within the imaging time, fewer BAK rings reached the maximum diameter of the mitochondrial fragment on which they were located compared to BAX rings [FigS1B], while the area of the mitochondrial fragments themselves remained largely constant [FigS1C]. However, the shapes of the mitochondria in the BAX– and BAK-overexpressing cells were different. To quantify the shape of the mitochondria we determined the aspect ratio (AR) between the shortest and longest axis of the mitochondrial fragments. Accordingly, a low AR corresponds to a still largely tubular mitochondrion, while an AR close to one corresponds to a completely rounded mitochondrial fragment [FigS1D]. We found that many of the mitochondrial fragments harboring BAK rings had a low AR of down to ∼ 0.2 at the start of ring formation, whereas mitochondrial fragments with BAX rings exhibited ARs of 0.6 or more at the start of ring formation. Almost all mitochondria with BAX rings reached an AR of close to 1, while the AR of many mitochondria with BAK rings remained significantly lower [Fig 1C]. This indicates that BAK is able to form rings on still elongated mitochondria, while BAX rings only occur on rounded mitochondrial fragments [FigS1D]. Furthermore, while in the Halo-BAX-overexpressing cells, we found exclusively one BAX-ring per mitochondrial fragment, in the Halo-BAK overexpressing cells, we sometimes found two BAK rings on one mitochondrion [FigS1E]. Another difference between the BAK– and BAX-overexpressing cells are the apoptotic ultrastructures formed by the two proteins. While BAX formed only clusters and rings on mitochondrial fragments in apoptotic cells, BAK additionally formed long, thin protrusions localized in areas between mitochondria, which we termed “linkers” [Fig 1B, arrowheads].

In summary, our live-cell STED data show distinct differences in the dynamics of oligomerization between overexpressed BAX and BAK in apoptotic cells.

### Endogenous BAX and BAK together form mosaic rings in cellulo

To investigate if overexpression of BAX and BAK induces BAX-BAK-structures that are different from those in apoptotic WT cells, we established a BAX-BAK immunofluorescence labeling approach suitable for STED microscopy. We treated U-2 OS WT cells with Actinomycin D and ABT-737 to induce apoptosis and labeled the cells after fixation with antibodies targeting BAX, BAK and TOM20 (as MOM marker). STED imaging revealed distinct BAX-BAK structures in apoptotic cells, which were not detectable in untreated (non-apoptotic) cells [Fig 2A-B]. The BAX-BAK structures localized predominantly to mitochondria (as shown by colocalization with TOM20), and we observed clusters, elongated shapes and rings. While some of the structures seemed to be devoid of one of the two proteins [Fig 2Ci, narrow and wide arrowheads], the vast majority of these structures, and especially the rings, were composed of both BAX and BAK [Fig 2Cii, arrows]. These results suggest that at endogenous expression levels, BAX and BAK together coalesce into apoptotic structures, and, in particular, apoptotic rings. A detailed examination of the apoptotic BAX-BAK structures of more than 150 cells revealed that virtually no “linkers” between mitochondria (not colocalizing with TOM20) are found in these cells. The large number of BAK “linkers” in the live-cell STED data [Fig 1B] is therefore apparently a consequence of overexpression.

**Fig 2.**
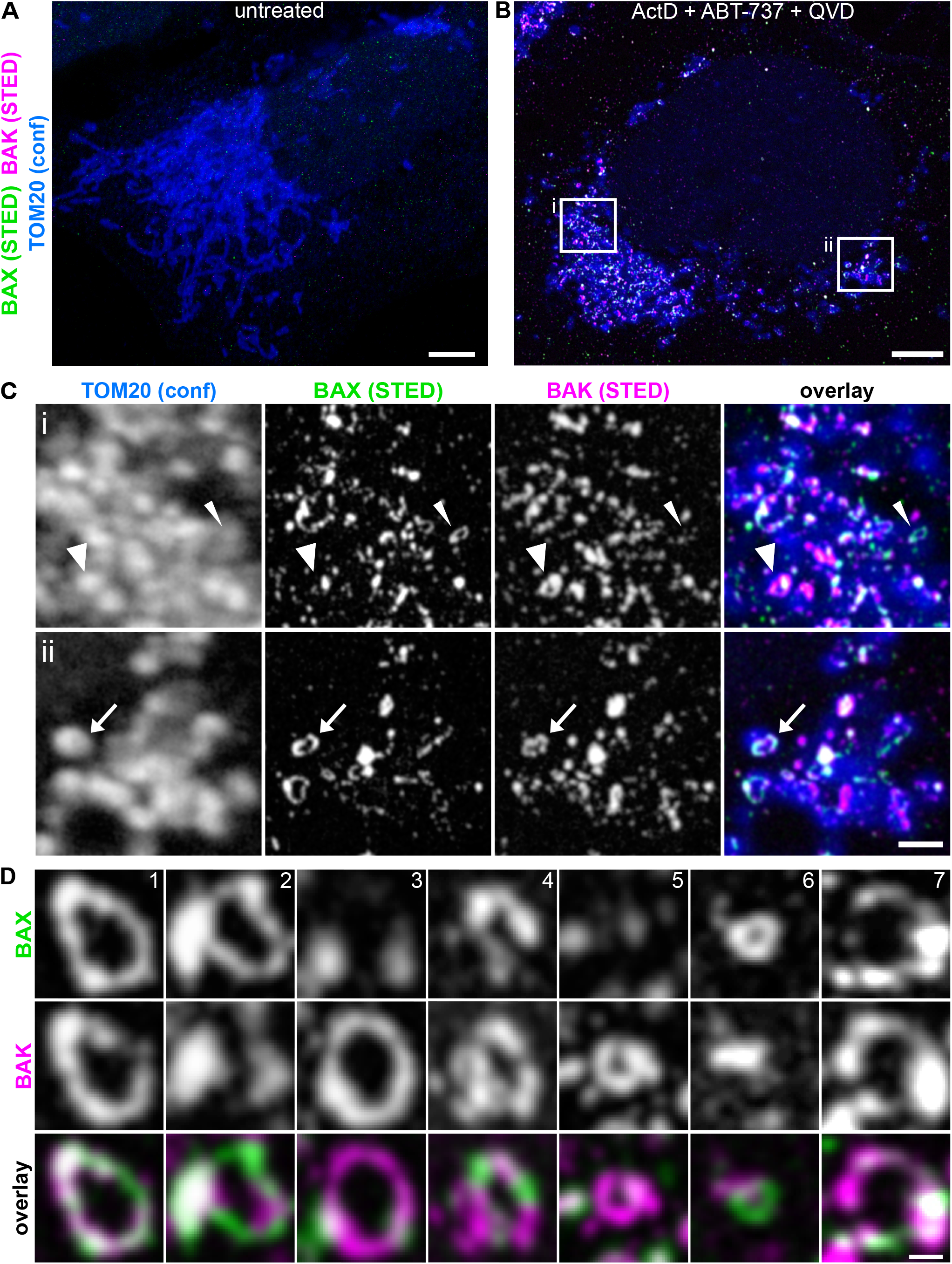
Endogenous BAX and BAK together form mosaic rings *in cellulo*. (**A**) STED image of an untreated, fixed U-2 OS WT cell, immunolabeled for endogenous BAX (green, STED), BAK (magenta, STED) and TOM20 (blue, confocal). (**B**) STED image of an apoptotic, fixed U-2 OS WT cell, immunolabeled for endogenous BAX (green, STED), BAK (magenta, STED) and TOM20 (blue, confocal). The cells were treated for 18h with 10 µM ABT-737, 10 µM Actinomycin D and 20 µM QVD-OPh. (**C**) Enlarged insets from (B). (i) Some apoptotic rings are largely comprised of either BAK (wide arrowhead) or BAX (narrow arrowhead). (ii) Most rings contain both BAX and BAK (arrow). (**D**) Exemplary images showing the great heterogeneity in size and composition of the BAX-BAK rings. Scale bars: 5 µm (A and B), 1 µm (C), 200 nm (D).

We also analyzed the distribution of BAX and BAK in apoptotic human primary cells. To this end, we induced apoptosis in human dermal fibroblasts (HDFa) and we found the same type of apoptotic structures, such as rings, composed of endogenous BAX and BAK [FigS2]. This shows that BAX-BAK structures and especially rings do not only occur in immortalized cancer cells but are a physiological phenomenon during apoptosis in human cells.

### Endogenous BAX and BAK are capable of forming apoptotic rings independently of each other

Qualitative evaluation of the apoptotic rings consisting of endogenous BAX and BAK showed that in some rings the fluorescence signals from the two proteins overlap [Fig 2D, example 1], while in other rings one of the two proteins dominates the ring outline [Fig 2D, example 2-3] or BAX and BAK alternate [Fig 2D, example 4]. The size of the rings is highly variable, likely due to different times of initiation of pore growth [Fig 2D, example 1-7]. In many rings, BAX and BAK showed an inhomogeneous distribution along the ring outline, forming large clusters in some places and being completely absent in other sections of the ring outline [Fig 2D, example 7].

The heterogeneity and the almost complete absence of one of the two protein partners in some rings suggested that BAX and BAK might be capable of forming rings independently from each other. It has been shown that BAX and BAK are largely redundant in their function to induce apoptosis (11, 12) and both BAX and BAK alone are capable of forming rings independently when overexpressed in a KO background (20, 21). However, the formation of apoptotic rings by endogenous BAK or BAK alone has not yet been demonstrated. Therefore, we generated a BAX-KO and a BAK-KO cell line in U-2 OS cells using CRISPR/Cas9 [Fig 3A] and found that both BAX as well as BAK are able to form rings in apoptotic cells independently of the other protein, even at endogenous expression levels [Fig 3B-C].

**Fig 3.**
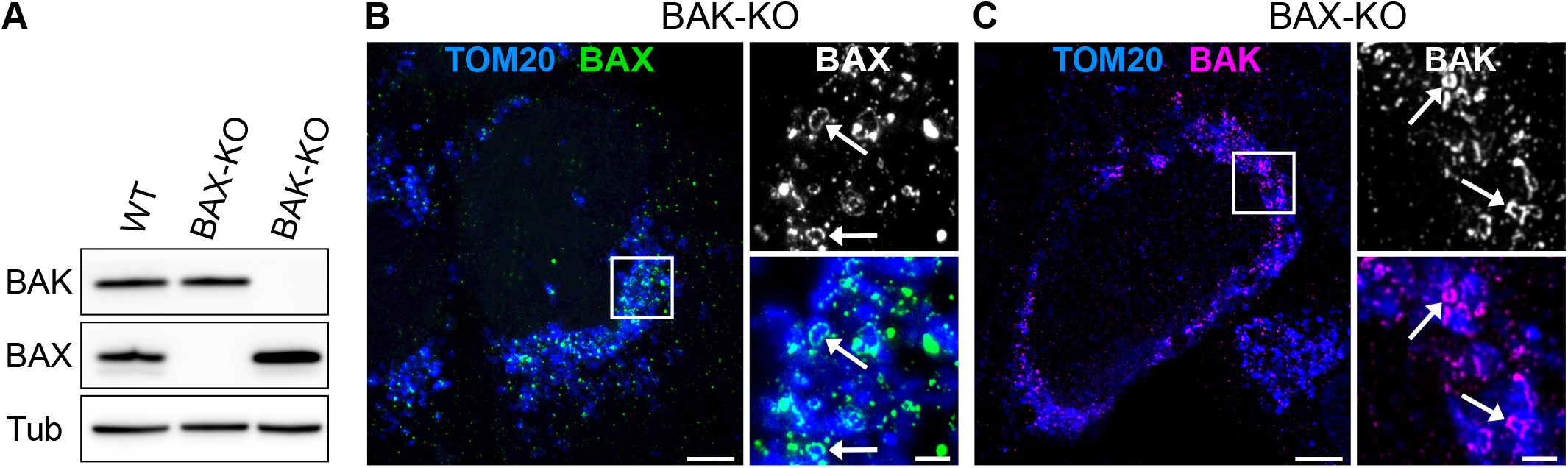
BAX and BAK are able to form rings and other apoptotic structures independently of each other. (**A**) Western blot analysis of CRISPR/Cas9-engineered BAX– and BAK-single-KO cell lines used in this study. (**B**) Left panel: STED image of fixed U-2 OS BAK-KO cells immunolabeled for endogenous BAX (green, STED), and TOM20 (blue, confocal). The cells were treated for 20h with 10µM Actinomycin D, 10µM ABT-737 and 20µM QVD-OPh. Right panels: Enlarged view of the BAX signal alone (top) or BAX signal and TOM20 signal (bottom) from the indicated area in the overview. Arrows indicate BAX rings. (**C**) Left panel: STED image of fixed U-2 OS BAX-KO cells immunolabeled for endogenous BAK (magenta, STED), and TOM20 (blue, confocal). The cells were treated for 20h with 10µM Actinomycin D, 10µM ABT-737 and 20µM QVD-OPh. Right panels: Enlarged view of the BAK signal alone (top) or BAK signal and TOM20 signal (bottom) from the indicated area in the overview. Arrows indicate BAK rings. Scale bars: 5 µm (B and C, left panel), 1 µm (B and C, right panels).

### BAX-BAK rings are heterogeneous in size and composition

We examined the rings in 3D using 4Pi-STORM microscopy, in order to explore if 2D STED microscopy is sufficiently accurate for quantification of apoptotic rings [FigS3]. 4Pi-STORM microscopy provides a localization precision in all three dimensions of 2-3 nm (27), at the expense of longer recording times. We found that about 75% of BAX rings were detectable in a single 2D plane extracted from the 4Pi-STORM image. We conclude that 2D STED microscopy, which enables the comparatively fast and robust acquisition of reliable data (28), is suitable for the analysis of rings on apoptotic mitochondria.

Thus, we set up an automated high-throughput 2D STED imaging workflow to analyze the size and composition of BAX-BAK rings quantitatively. In the resulting images, we manually traced the outlines of 530 rings [Fig 4A]. We then plotted the fluorescence intensities of the BAX and BAK channels along the ring outline [Fig 4B]. The circumference of the rings ranged from 0.43 µm to 3.81 µm with a median of 1.21 µm [Fig 4C]. These ring circumferences correspond to circle diameters of 140 to 1210 nm with a median of 380 nm, which is consistent with published diameters of endogenous BAX rings (19) and our live-cell STED data. The values are slightly larger than previously measured diameters of overexpressed BAX or BAK rings (20, 21).

**Fig 4.**
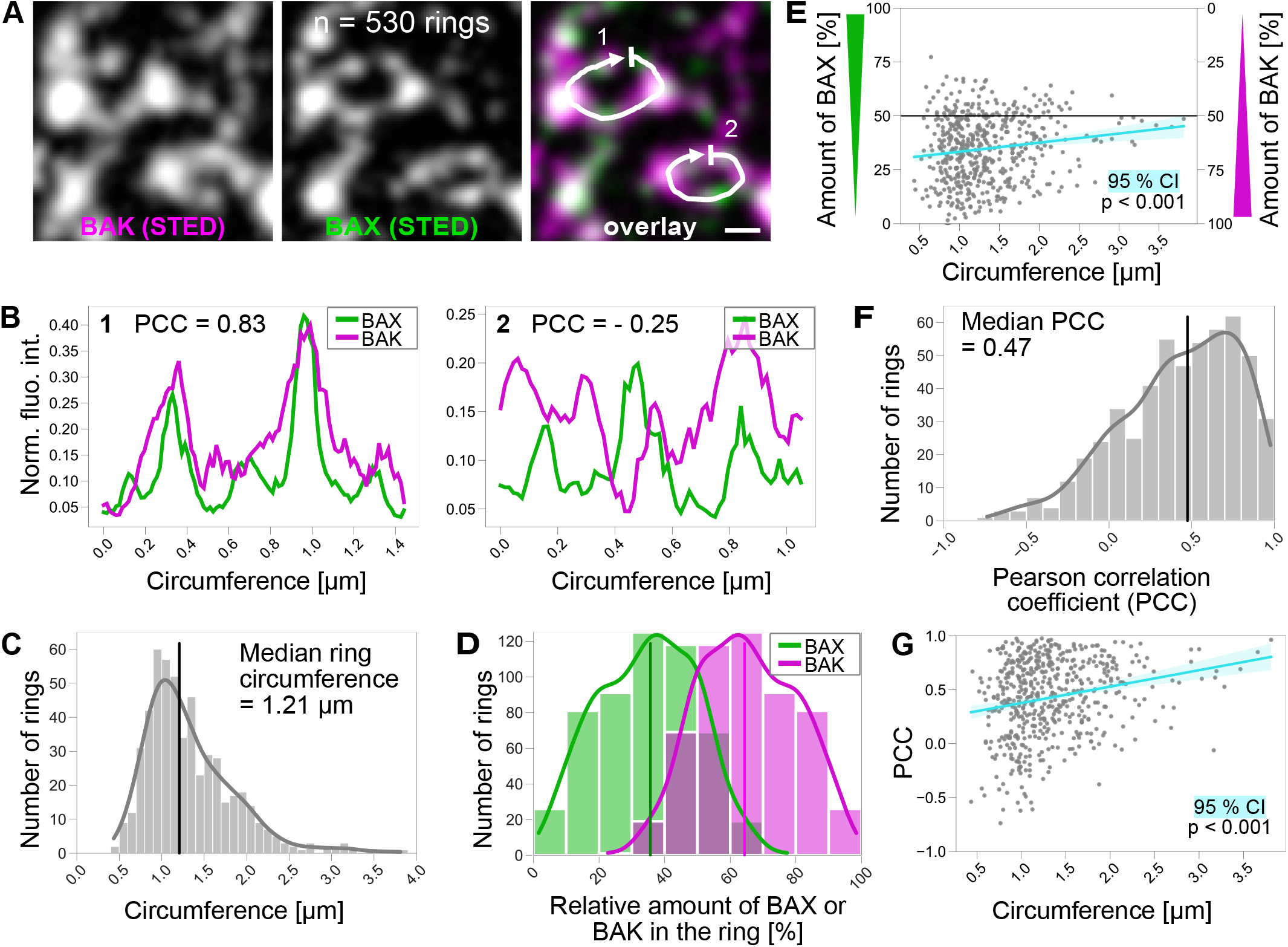
BAX-BAK rings are heterogeneous in size and composition. (**A**) Exemplary STED image of BAX-BAK rings with manually drawn line profiles. (**B**) Fluorescence intensity plots of ring line profiles (same as A). The normalized fluorescence intensity of the BAX (green) and BAK (magenta) channels was plotted (Norm. fluo. int.) along the circumference of the ring. PCC = Pearson correlation coefficient. (**C**) Circumferences of all BAX-BAK rings ranged from 0.43 to 3.81 µm. Vertical black line indicates the median of 1.21 µm. Bin width = 100 nm. (**D**) Relative amounts of ring outline occupied by BAX (green) and BAK (magenta) in all rings. Vertical lines indicate the medians of 36 % for BAX and 64 % for BAK. Bin width = 10 %. (**E**) Relative amounts of BAX (left y-axis) and BAK (right y-axis) as a function of the ring size (circumference). The black horizontal line at 50% represents equal distribution of BAX and BAK. The turquoise line shows the linear regression of the data and light turquoise area shows the 95 % CI with p < 0.001. (**F**) Pearson correlation coefficients (PCC) of BAX and BAK in the rings. The PCCs in the rings range from – 0.74 to 0.98 with a median at 0.47 (vertical black line). Bin width = 0.1. (**G**) PCC as a function of the ring size (circumference). The turquoise line shows the linear regression of the data and the light turquoise area shows the 95 % CI with a p < 0.001. Scale bar: 200 nm (A). n = 530 rings from three independent experiments.

To determine the composition of the rings, we examined the relative amounts of BAX and BAK covering the ring outline. By comparing the fluorescence intensity values of the BAX and BAK channels in the 530 rings, we found that most rings contained both BAX and BAK, but that BAK was generally more abundant. The median relative proportion of BAK and BAX signals in the apoptotic rings was 64% and 36%, respectively [Fig 4D]. We found that with increasing ring size, the relative amounts of BAX and BAK in the rings approached 50% [Fig 4E]. This suggests that BAX and BAK are more likely to be present in equal proportions in larger rings.

To analyze the spatial distribution of BAX and BAK relative to each other along the ring outline, we calculated the Pearson correlation coefficient (PCC) of the fluorescence intensity profiles. We found a PCC distribution from –0.74 to 0.98 with a median of 0.47, representing an overall positively correlated localization of BAX and BAK in the rings [Fig 4F]. The positive correlation indicates that BAX and BAK overlap to some extent along the ring outlines in most rings, suggesting that BAX and BAK (at the given resolution of the STED microscope of around 40 nm) coalesce into the same spots and are not mutually exclusive (within this size scale) along the ring outline. The PCC also increased with increasing ring sizes [Fig 4G], which fits well with the increasing equilibration of the amounts of BAX and BAK in larger rings.

In summary, we have shown that overexpression of BCL-2 effectors can induce ultrastructures that are not obvious at endogenous protein levels. Endogenous BAX and BAK form mainly MOM-associated structures, specifically rings, also in non-immortalized cells. These rings are lining the apoptotic pores as unordered, mosaic arrangements, and each protein alone is sufficient to form these apoptotic rings. Together with the live-cell STED data, which showed that rings grow over time, our results suggest a temporal hierarchy in the accumulation of BAK over BAX during ring formation. BAK accumulates in the rings before BAX, and in the growing rings both BAK and BAX are recruited until they reach approximately equal amounts in larger rings.

## Discussion

In this study, we used STED super-resolution microscopy to uncover the dynamics and composition of BAX and BAK in apoptotic pores *in cellulo*. We found differences in pore formation at the nanoscale, depending on the protein. We observed that BAK is able to form pores on still elongated mitochondria while BAX pores were only found on rounded up mitochondrial fragments [Fig 1C]. This difference might result from the direct interaction of BAX with the mitochondrial fission protein Drp1 (29), or the fact that BAX oligomerization induces faster depolarization of the mitochondria. Furthermore, we found that BAX rings grow slightly larger than BAK rings, both in absolute size and in relation to the mitochondrial fragments on which they are located [FigS1A-B]. Somewhat unexpectedly, in BAK overexpressing cells, we observed many BAK-protrusions that are not located on mitochondria [Fig 1B]. These “linkers” were virtually absent in non-overexpressing apoptotic cells in which endogenous BAK was labeled, suggesting changes in BAK oligomerization due to overexpression. We conclude that studies of the oligomerized ultrastructures of BCL-2 effectors should be confirmed by investigating these proteins at endogenous expression levels.

Generally, BAX-BAK rings were not smooth, perfect rings, but exhibited irregularities such as gaps in the BAX/BAK signal or larger BAX/BAK clusters associated to it. The analysis of the distribution of endogenous BAX and BAK in the apoptotic rings did not reveal a particularly ordered arrangement of the two proteins. BAX and BAK colocalize to different extents in some parts of some of the rings and do not colocalize in other parts [Fig 2D, Fig 4A-B, F]. Thus, BAX and BAK do not seem to form an ordered wall of single-layer BAX or BAK dimers around the pore edge, but rather an unordered arrangement with voids and large BAX/BAK clusters on the ring outline.

The BAX-BAK rings were not only heterogeneous with respect to size, shape and regularity, but also heterogeneous in their BAX/BAK content. Although the relative amount of BAX and BAK varied greatly from one ring to another, BAK was generally more abundant in small rings [Fig 4D]. The amount of BAX in the apoptotic rings increased with increasing ring size, resulting in BAX and BAK being equally abundant in most large rings [Fig 4E]. The prevalence of BAK, especially in smaller rings, could be because BAK is already present at the MOM in non-apoptotic cells. This allows BAK to form clusters and then rings earlier, while BAX is still recruited from the cytosol to the MOM (14). This is consistent with our observation that BAK can already form rings on still elongated mitochondria [Fig 1B-C]. It is furthermore in line with previous findings that BAK but not BAX has a high tendency to autoactivate (30) and that BAK oligomerizes faster into apoptotic structures (21). Thus, smaller rings with little BAX are likely to represent early intermediates. Taken together, our data suggest a temporal hierarchy in the oligomerization of BAK over BAX during apoptotic ring formation.

In summary, we show that BAX and BAK together form mosaic rings on apoptotic mitochondria. BAX and BAK are recruited differently into the rings, with BAK temporally preceding BAX and the distribution of the two proteins in the rings is in line with the model of a toroidal apoptotic pore.

## Acknowledgements

We thank Peter Ilgen for help with illustrations, Till Stephan and Christian Brueser for fruitful discussions, Joerg Bewersdorf for providing us with the OMP25 expression plasmids and the Facility for Synthetic Chemistry at the Max Planck Institute for Multidisciplinary Sciences for synthesizing live-cell dyes. This work was supported by the European Research Council (ERCAdG No. 835102) and by the DFG-funded SFB 1456 – project C06 (both to S.J.).

## Author contributions

S.V.S and S.J. designed research. S.J. conceived the project. S.V.S., A.F and M.B. performed research. S.V.S and J.K.-F. analyzed data. S.V.S wrote the initial draft and S.V.S, D.C.J. and S.J. wrote the final paper. All authors approved the manuscript.

## Materials and methods

### Antibodies

|  | Supplier | Cat# |
| --- | --- | --- |
| *Immunofluorescence antibodies* |  |  |
| Mouse anti human BAX Monoclonal Antibody (2D2) | ThermoFisher/invitrogen | # MA5-13994 |
| Rabbit anti human BAX antibody, clone 1C7, ZooMAb® Rabbit Monoclonal | Merck/Sigma-Aldrich | # ZRB1103-4X25UL |
| Rabbit anti human BAK Recombinant Monoclonal Antibody (SU32-07) | ThermoFisher/invitrogen | # MA5-32111 |
| Mouse anti human BAK antibody monoclonal [AT38E2] | abcam | # ab104124 |
| Mouse anti human TOM20, purified, monoclonal, Clone 29 | BD Transduction Laboratories | # 612278 |
| Rabbit anti human TOMM20 antibody [EPR15581-39], recombinant, monoclonal | abcam | # ab186734 |
| Mouse IgG anti-Chicken IgY (H+L)-unconj., MinX none | dianova | # SBA-8320-01 |
| Rabbit Anti-Chicken IgY H&L | abcam | # ab97136 |
| Rabbit anti human TOMM20 antibody [EPR15581-39], coupled to Alexa Fluor® 488 | BD | # 560263 |
| *Western blot antibodies* |  |  |
| Rabbit anti human monoclonal recombinant Anti-BAX antibody [EPR18284] | abcam | # ab182734 |
| Rabbit anti human polyclonal Anti-BAK Antibody | Merck/Millipore | # 06-536 |
| Goat Anti-Rabbit Peroxidase-AffiniPure IgG (H+L), Conjugation: Horseradish Peroxidase (HRP) | Dianova | # 111-035-144 |
| Goat Anti-Mouse Peroxidase-AffiniPure IgG (H+L), Conjugation: Horseradish Peroxidase (HRP) | Dianova | # 115-035-062 |
| Mouse anti-α-Tubulin antibody, monoclonal, clone B-5-1-2I | Sigma | # T6074 |

### Cloning of plasmids

#### Overexpression plasmids

##### Halo-BAX

hBAX C3-EGFP was a gift from Richard Youle (Addgene plasmid # 19741). The plasmid was linearized using the restriction endonucleases AgeI and SacI. The Halo tag was amplified by PCR from plasmid pENTR4-HaloTag (w876-1), which was a gift from Eric Campeau (Addgene plasmid # 29644), with the following primers: Halo_fw_AgeI: 5’-gctaccggtcgccaccatggcagaaatcggtac-3’ Halo_rv_SacI: 5’-tgagctcgagatctgagtagccggaaatctcgagcgtcgaca-3’ The amplified Halo-tag was then inserted into the linearized BAX plasmid.

##### Halo-BAK

EGFP-BAK was a gift from Richard Youle (Addgene plasmid # 32564). The plasmid was linearized using the restriction endonucleases AgeI and XhoI. The Halo tag was amplified by PCR from plasmid pENTR4-HaloTag (w876-1), which was a gift from Eric Campeau (Addgene plasmid # 29644), with the following primers: Gibson_fw_Halo-BAK_AgeI: 5’-gatccgctagcgctaccggtatggcagaaatcggtactggc-3’ Gibson_rv_Halo-BAK_XhoI: 5’-agccataagcttgagctctagatctgagtagccggaaatctcgagc– 3’ The amplified Halo-tag was then inserted into the linearized BAK plasmid via Gibson assembly.

#### Plasmids for OMP25 – safe harbor locus (AAVS1) integration

##### pX330-AAVS1

The gRNA+Cas9 plasmid pX330-AAVS1 to cut the safe harbor locus was derived from pX330-U6-Chimeric_BB-CBh-hSpCas9, which was a gift from Feng Zhang (Addgene plasmid # 42230). The following oligonucleotides were annealed and integrated into pX330 after linearization with the BbsI restriction endonuclease:

AAVS1-gRNA-fw: 5‘-CACCGtgtccctagtggccccactg-3‘

AAVS1-gRNA-rev: 5‘-AAACCAGTGGGGCCACTAGGGACAC-3‘

##### SNAP-OMP25 and GFP-OMP25

To generate the donor plasmid to integrate SNAP-OMP25 or GFP-OMP25 into the safe harbor locus (AAVS1), the plasmids AAVS1-Snap-OMP25 and AAVS1-GFP-OMP25 were created. For this, AAVS1-Basticidin-CAG-Flpe-ERT2, which was a gift from Su-Chun Zhang (Addgene plasmid #68461), was linearized by using the restriction endonucleases SalI and EcoRV.

GFP-OMP25 and Snap-OMP25 were gifts from the Bewersdorf lab (Yale University) and are described elsewhere (31, 32). OMP25 with the respective tag was amplified by PCR using the primers given below and subsequently integrated into the linearized AAVS1 plasmid by Gibson assembly.

Gibson_AAVS1_Snap-OMP25_fw:

5’-tcattttggcaaagaattcgtcgacgccgccaccATGGACAAAGACTGCGAAAT-3’

Gibson_Snap-OMP25_AAVS1_rv:

5’-tgattatcgataagcttgatatcTCAAAGTTGTTGCCGGTATC-3’

Gibson_AAVS1_GFP-OMP25_fw:

5’-tcattttggcaaagaattcgtcgacGCCGCCACCATGGTGAGC-3’ Gibson_GFP-OMP25_AAVS1_rv:

5’-gttgattatcgataagcttgatatctcaaagttgttgccggtatctcatg-3’

#### Plasmids for BAX and BAK knockout (KO)

##### pX458-BAX and pX458-BAK

The gRNA+Cas9 plasmids pX458-BAX or pX458-BAK to cut the first exon of the genes was derived from pSpCas9(BB)-2A-GFP (PX458), which was a gift from Feng Zhang (Addgene plasmid # 48138). The following oligonucleotides were annealed and integrated into pX458 after linearization with the BbsI restriction endonuclease:

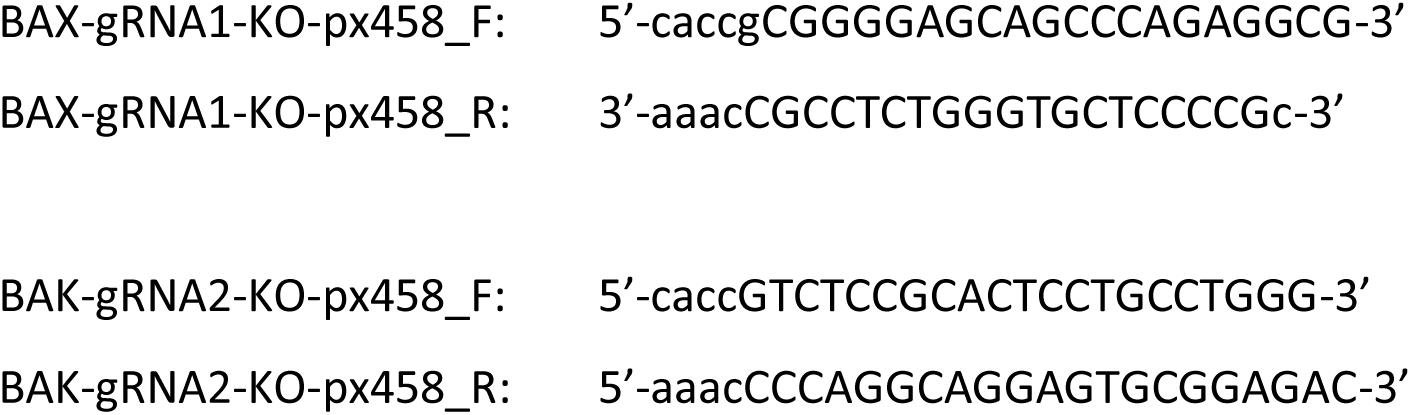

##### Cell culture

U-2 OS cells (HTB-96™, ATCC) were cultured in McCoy’s 5A medium with 10% FBS (Gibco or Sigma) and 1mM sodium pyruvate. HDFa (PCS-201-012™, ATCC) were cultured in Dulbecco’s Modified Eagle Medium (DMEM) with 4.5 g/L glucose, GlutaMax™, 1 mM sodium pyruvate and 10% FBS. Bacterial contamination of the cell lines was suppressed by addition of 100 µg/mL streptomycin and 100 U/mL penicillin to the culture medium. The cells were grown in a CO_2_ incubator at 37 °C, 5% CO2 and 90% humidity. Before the cells reached confluency, they were split by trypsinization up to a 1:10 dilution and discarded upon reaching p20.

### Generation of cell lines with stable safe harbor locus (AAVS1) integration via CRISPR/Cas9

To generate a stable cell line expressing tagged OMP25 in the safe harbor locus (AAVS1), U-2 OS cells were co-transfected with the plasmids AAVS1-Snap-OMP25 or AAVS1-GFP-OMP25 and pX330-AAVS1 by electroporation with the Amaxa cell line nucleofector. Starting two days after transfection, the cells were selected using McCoy’s medium containing 2.5 – 5 µg/mL Blasticidin S (Invivogen # ant-bl-05) for 7 days. Two weeks after transfection, the Snap-OMP25 cells were stained with McCoy’s medium containing 500 nM SiR-BG (synthesized by the Facility for Synthetic Chemistry, Max-Planck Institute for Multidisciplinary Sciences) for 1 hour. GFP-OMP25 containing cells were not stained. Using fluorescence-activated cell sorting (FACS), single fluorescent cells were transferred into 96 well plates. After about 3 weeks, single cell clones were selected under the microscope for bright and correctly localized fluorescence. Proper integration of the OMP25 fusion protein was verified by PCR with the following primers:

AAVS1_genom_wt_fw: 5‘-CCCCTATGTCCACTTCAGGA-3’

AAVS1_genom_wt_rv: 5‘-CAGCTCAGGTTCTGGGAGAG-3’

AAVS1_genom_intg_fw: 5‘-GTCGTGCCAGCGGATCGACAGTA-3’

AAVS1_genom_intg_rv: 5‘-GGAGAAGGATGCAGGACGAGAAACACAGC-3’

### Generation of BAX KO and BAK KO cell lines via CRISPR/Cas9

To generate cell lines with knocked out BAX or BAK, U-2 OS cells were transfected with 2 µg of the plasmids pX458-BAX or pX458-BAK by electroporation with the Amaxa cell line nucleofector. Three days after transfection, single cells were transferred into 96 well plates via fluorescence-activated cell sorting (FACS) for GFP. After about 3 weeks, single cell clones were clonally expanded and checked for the absence of the BAX or BAK proteins by Western Blot. Selected clones were then analyzed by next generation sequencing (Illumina) of the targeted exon.

### Preparation of cells for live-cell imaging with Halo-BAX or Halo-BAK

U 2-OS cells stably expressing Snap-OMP25 or GFP-OMP25 were grown to 50-70%-confluency in a 35-mm culture dish (ibidi GmbH # 81158) and transfected with 1 µg of the Halo-BAX or Halo-BAK plasmid in Lipofectamine® 2000 Reagent (ThermoFisher Cat#11668019). After 2h of expression the medium was changed and medium containing 20 µM QVD was added, in order to prevent the detachment of apoptotic cells. 4h after transfection some cells started to undergo apoptosis due to the expression of BAX or BAK. Live-cell movie recording was started 4-12 hours after transfection. For this, the cells were incubated with medium containing 500 nM SiR-BG (to label Snap-tag, if Snap-OMP25 was present) and 250 nM Atto590-CA (to label Halo-tag) (both synthesized by the Facility for Synthetic Chemistry, Max-Planck Institute for Multidisciplinary Sciences) for one hour (32). To washout the unbound dyes, the cells were incubated in medium without dye for one hour. Before imaging, the medium was changed to Fluorobrite (ThermoFisher #A1896701) with 10% FBS. All steps after transfection were performed in the presence of 20µM QVD.

### Apoptosis induction with Actinomycin D

Where indicated, apoptosis was induced using Actinomycin D and ABT-737. 10µM Actinomycin D (Merck # 114666-5MG), 10 µM ABT-737 (ApexBio #A8193) and 20 µM QVD (ApexBio #A1901) were added to the cells in full medium for 10-20h before fixation.

### Immunofluorescence staining

Cells were grown on 12 mm #1.5H glass coverslips (Marienfeld) overnight. Then apoptosis was induced (see above) where indicated. Cells were fixed in 8% FA/PBS (Thermo Scientific™ #28908) for 10 min, followed by permeabilization in 0.5% Triton-X-100/PBS (Merck/Millipore #1086031000) for 5 min. Cells were blocked for 1 h in 5% BSA/PBS (Albumin (BSA) Fraction V, (pH 7.0), AppliChem # A13910500) and incubated with primary antibodies in 5% BSA/PBS for 1h at room temperature in a humidified chamber. The cells were then washed in PBS three times and secondary antibodies in 5% BSA/PBS were added for 1 h at room temperature in a humidified chamber. Samples were washed 3x in PBS and mounted in Prolong Diamond mounting media with or without DAPI (ThermoFisher # P36966 or #P36965).

If an antibody directly coupled to a fluorophore (TOM20-Alexa Fluor® 488, from here on called tertiary antibody) was included, the samples were incubated for 1h with a blocking antibody raised in the same species as the tertiary antibody in order to block unbound epitopes of the secondary antibody. Then the sample was washed 3x in PBS and the tertiary antibody was added in BSA/PBS for 1h at room temperature in a humidified chamber. After washing 3x in PBS, the samples were mounted as described above.

### Western blot

To generate whole cell protein lysates, cells from a confluent 6-well plate were harvested by trypsinization and spinning at 300 g for 5 min. The pellet was resuspended in lysis buffer [50 mM Tris-HCl pH7.4, 4 mM MgCl2, 0,1 mM DTT, 1% SDS; prepare 1000 µL, then add fresh: 50 µL cOmplete solution (cOmplete, EDTA-free Protease inhibitor cocktail tablet, Merck # 5056489001; 25X stock solution prepared according to manufacturer’s instructions) and 2 µL Benzonase (Sigma/Merck # E1014)]. The lysates were mixed with 6x sample buffer [0.05 M Tris-HCl pH6.8, 1 % SDS, 1 % ß-Mercaptoethanol, 10 % Glycerin, 0.001 % Bromphenolblue] and heated for 5 min at 95 °C.

NuPage Novex 4-12% BT Midi gels (Thermo Fisher #STM4003) were placed into an XCell4 SureLock™ Midi-Cell gel tray (ThermoFisher #WR0100) and filled with 1X NuPAGE™ MES SDS Running Buffer (20x) (Invitrogen/ThermoFisher #NP0002). Lysates and 5 µL of ladder (PageRuler™ Prestained Protein Ladder, 10 to 180 kDa Therm Scientific™ # 26617) were loaded into the gel pockets. The gels were then run at 150V for 80 min.

After running, the gels were blotted onto nitrocellulose membranes (iBlot stack, invitrogen # IB23002) by using the iBlot 2 system (invitrogen #IB21001) according to the manufacturer’s instructions. The blotted membrane was immersed in block buffer [5% milk (AppliChem # A0830.1000) in 1XTBS (TBS, Thermo Fisher #28358)] for 2 hr at RT. The block buffer was discarded and the membrane was washed 1 x 15 min with TBS-T (Thermo Fisher #28360). The primary AB, diluted in block buffer was added. The membrane with primary AB solution was incubated o/n at 4 °C. The next day the membrane was washed 3 x 5 min with TBS-T on RT shaking. Then the secondary AB coupled to HRP was diluted freshly in block buffer and added to the membrane, which was incubated at RT for 1 hr shaking and shielded from light. The membrane was washed 3 x 5 min with TBS-T on RT shaking. After a brief rinse in TBS, ECL solution (Immobilon Forte Western HRP substrate, Millipore #WBLUF0100) was used to develop the membrane, which was imaged on an Amersham™ Imager 600.

### STED imaging

Live-cell STED videos were acquired on a STED microscope (Expert Line, Abberior Instruments) equipped with a UPlanSApo 100x/1,40NA oil objective (Olympus). 488, 561 and 640 nm excitation lasers and a 775 nm depletion laser were used. The pinhole was set to 0.9 Airy units and a pixel size of 15-20 nm with a pixel dwell time of 5-10 µsec was used.

The fixed-cell STED recordings were acquired on a STEDycon (Abberior Instruments) mounted on a Nikon inverted microscope (Ti-2) with a CFI PlanApo 100x/1.45NA oil objective (Nikon). 488, 561, 640 nm excitation lasers and 775 nm depletion laser were used. The pinhole size was set to 1.13 Airy units and a pixel size of 15 nm with a pixel dwell time of 10 µsec and three line averages was used. Nikon NIS-elements software allowed multi-position acquisition and the integrated coding language was used to program a pipeline to communicate with the STEDycon and automatically acquire STED images at the predefined positions.

### 4Pi-STORM imaging

The samples for 4Pi-STORM imaging were prepared as described in (27). In brief, U 2-OS cells were seeded on custom-made coverslips, where one-quarter of the glass was covered by a thin aluminum layer to create a mirror surface needed for alignment of the sample and microscope objectives. The cells were treated to induce apoptosis (see above) or left untreated, fixed and stained with primary antibody (see immunofluorescence above). The primary antibody was detected by a Fab fragment coupled to Alexa Fluor 647 (Thermo Fisher, A21246; dilution 1:500). The sample was then covered in imaging buffer (27) and a second coverslip was placed on top. This sandwich of coverslips (containing the cell layer in between) was sealed and mounted vertically on the stage of the microscope. When a region of interest (ROI) was found and brought into focus, all Alexa Fluor 647 fluorophores in the ROI were switched off with high red laser intensities (642 nm at 10-20 kW/cm^2^). Stochastic fluorescence blinking events were then recorded on the camera. During the measurement, the ultraviolet laser (405 nm) was switched on, and its intensity slowly increased, to maintain a constant rate of fluorophore switching events over time. A camera exposure time of 10 ms was used and 100000 frames were recorded for each sample. STORM data analysis and image reconstruction was carried out using custom software (27). For visualization purposes, the 4Pi-STORM data was rendered using Blender, and quantification of rings detected in 2D vs. 3D was performed manually.

### Image processing

Only for illustration purposes, all images were contrast enhanced and the background was subtracted with a rolling ball algorithm in the BAX and BAK channels. The channels were then filtered with a Gaussian kernel with sigma of 1 or 2 pixels.

### Quantification and data analysis

All quantifications and data analyses were carried out on raw data.

#### Live-cell data

The diameter of the apoptotic rings and the lengths of the mitochondrial axes in the live-cell STED movies were measured manually in each frame using FIJI (33). The AR of the mitochondrial axes was calculated as the shortest mitochondrial axis length divided by the longest mitochondrial axis.

#### Analysis of STED images of endogenous BAX and BAK

The apoptotic rings were annotated in FIJI by manually tracing the ring outlines. The fluorescence intensity profiles were determined along the ring contour with a line width of 5 pixels. The subsequent analysis was performed in Python.

In brief: The ring circumference was determined. The BAX/BAK content was calculated by normalizing the line profiles to the maximum and mean value within the respective channel and replicate. Then the predominant channel was identified at each position. From this, the relative proportions of BAX and BAK along the ring outline were determined. The Pearson correlation coefficient (PCC) between the BAX and BAK intensity series along the ring contour was calculated. These data were plotted against each other and, where indicated, the linear regression together with the 95% confidence interval (CI) and the p-value was determined. The histograms were plotted with the indicated bin widths and a kernel density estimation with a Gaussian kernel was performed and plotted. The data for the analyses come from three independent replicates.

