## Supplemental Information for "Endogenous BAX and BAK form mosaic rings of variable size and composition on apoptotic mitochondria"

Sarah Vanessa Schweighofer et al.

**FigS1. Differences in BAX and BAK pore formation.**

- **(A)** Absolute ring diameter of BAX (green) or BAK (magenta) rings plotted over time.
- **(B)** Relative ring diameter (to longest mitochondrial axis) of BAX (green) or BAK (magenta) rings plotted over time.
- **(C)** Mitochondrial area of the mitochondrial fragments harboring BAX (green) or BAK (magenta) rings plotted over time.
- **(D)** Schematic illustration of AR (i) and relative ring diameter (ii).
- **(E)** Selected frames of a live-cell STED movie from a U-2 OS WT cell undergoing apoptosis. The cells stably overexpressed GFP-OMP25 (blue, confocal imaging mode) and were transiently transfected with Halo-BAK (labeled with Atto590, magenta). Cells were imaged in the presence of 20 µM QVD-OPh. Two BAK rings form on one mitochondrial fragment (arrows).

Data are quantified from 23 individual rings in 3 biological replicates of 2 independent experiments per condition.

Scale bar: 1 µm (E).

**FigS2. BAX-BAK rings in human dermal fibroblasts.**

**(A)** STED image of an apoptotic, fixed HDFa WT cell, immunolabeled for endogenous BAX (green, STED), BAK (magenta, STED) and TOM20 (blue, confocal). The cells were treated for 10h with 10 µM ABT-737, 10 µM Actinomycin D and 20 µM QVD-OPh.

**(B)** Enlarged insets from (A) showing mosaic BAX-BAK rings (arrow).

Scale bars: 5 µm (A), 1 µm (B).

**FigS3. BAX rings are located in the cell in all spatial orientations.**

**(A)** 4Pi-STORM image reconstruction of BAX in apoptotic cells. Data is displayed as perspective view along the z-axis with depth color coding. U-2 OS cells were treated with 10 µM ActD. 20 µM Q-VD-OPh was added to prevent the detachment of the cells from the coverslips. The apoptotic cells were fixed and labeled with the 2D2-BAX antibody, detected by a secondary Fab fragment coupled to Alexa Fluor 647, and prepared for 4Pi-STORM imaging. The image is a representative example of 3 replicates.

**(B)** Enlargement of the box in (A), as viewed from two angles turned by 90°, the first one showing only a line of BAX (left panel) which actually is a BAX ring when viewed from the side (right panel).

**(C)** Quantification of rings detected in 2D (gray area) vs 3D (full circle). Rings were counted in 5 datasets.

Scale bars: 2 µm (A), 250 nm (B).

### FigS1

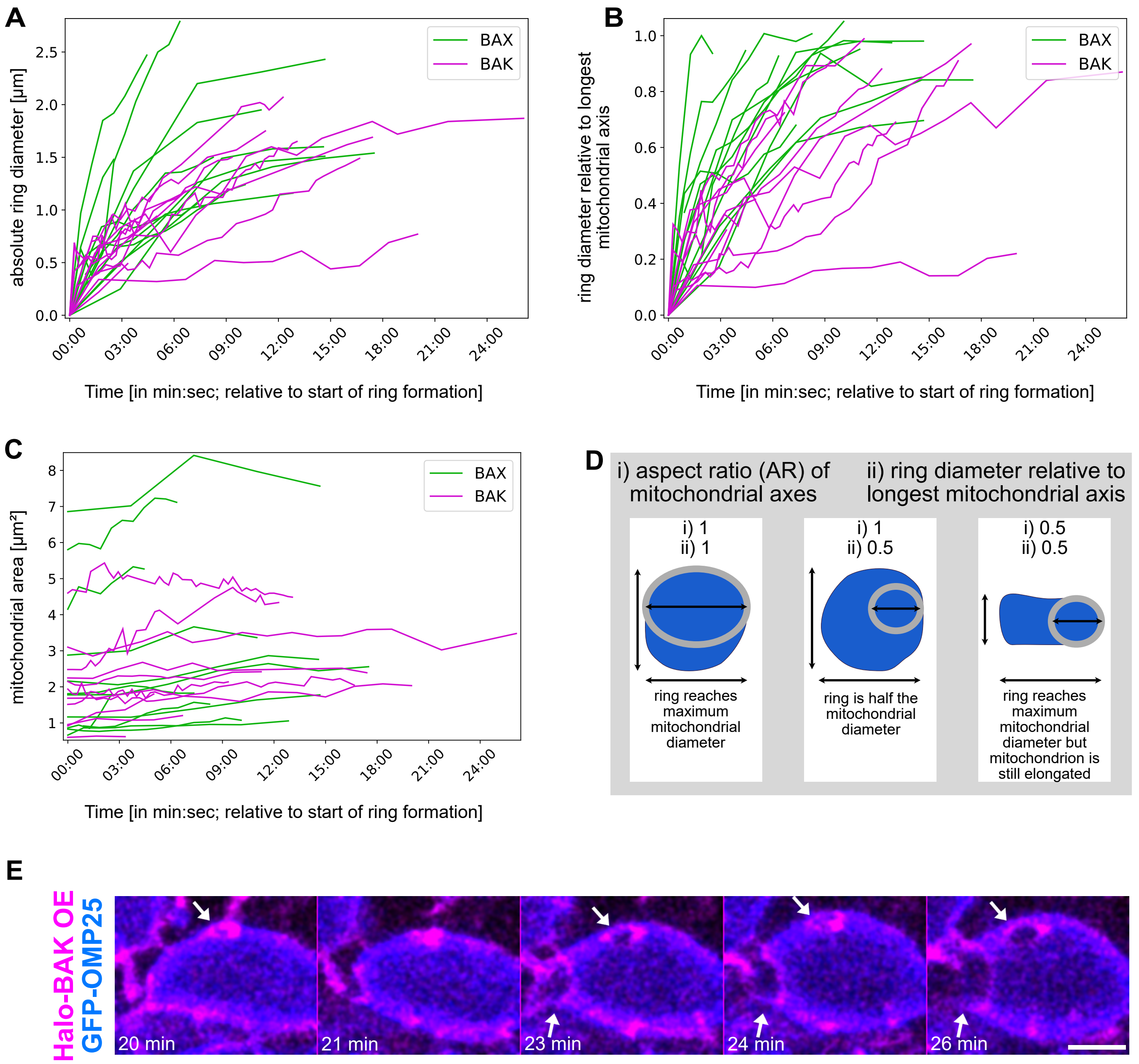

A
absolute ring diameter [µm]
Time [in min:sec; relative to start of ring formation]
BAX
BAK
B
ring diameter relative to longest mitochondrial axis
Time [in min:sec; relative to start of ring formation]
BAX
BAK
C
mitochondrial area [µm²]
Time [in min:sec; relative to start of ring formation]
BAX
BAK
D
i) aspect ratio (AR) of mitochondrial axes
ii) ring diameter relative to longest mitochondrial axis
i) 1
ii) 1
ring reaches maximum mitochondrial diameter
i) 1
ii) 0.5
ring is half the mitochondrial diameter
i) 0.5
ii) 0.5
ring reaches maximum mitochondrial diameter but mitochondrion is still elongated
E
Halo-BAK OE
GFP-OMP25
20 min
21 min
23 min
24 min
26 min

FigS2

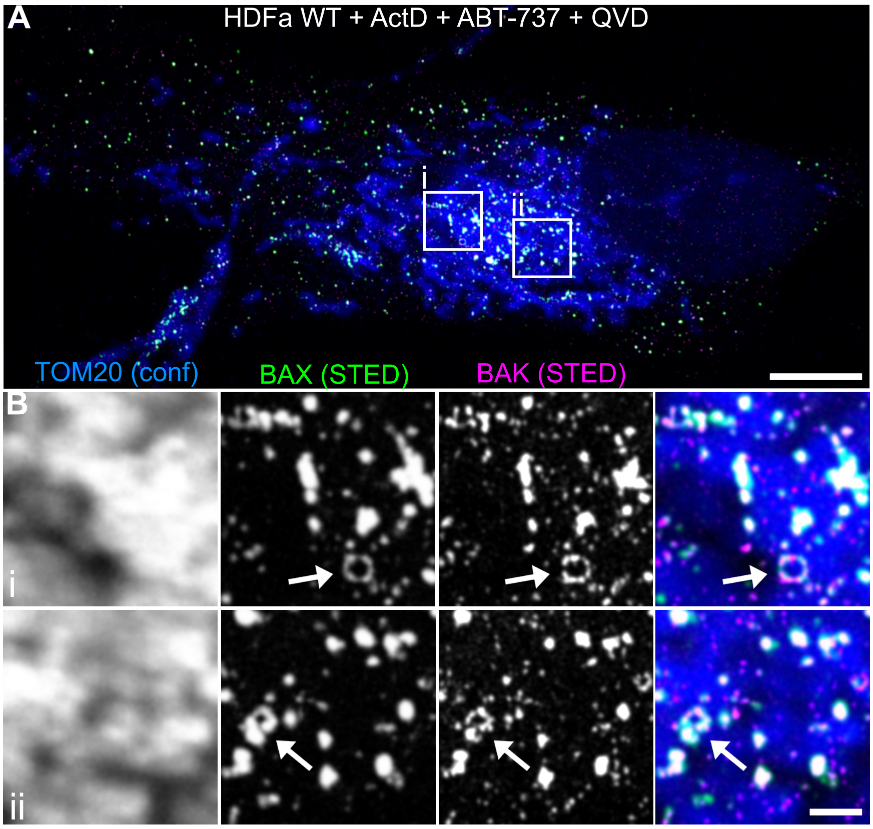
A
HDFa WT + ActD + ABT-737 + QVD
i
ii
TOM20 (conf)
BAX (STED)
BAK (STED)
B
i
ii

FigS3

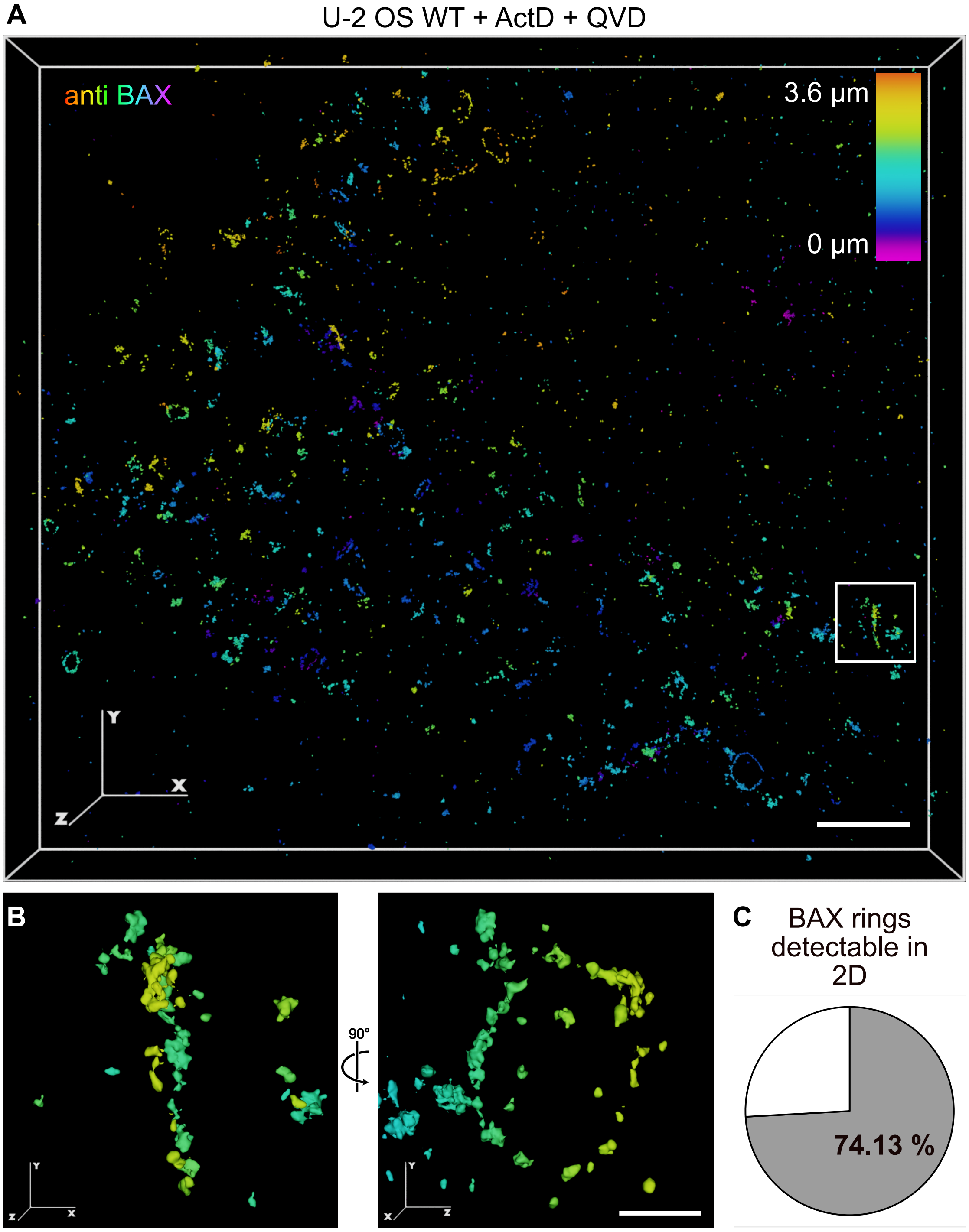
A
U-2 OS WT + ActD + QVD
anti BAX
3.6 µm
0 µm
Y
X
Z
B
90°
C
BAX rings detectable in 2D
74.13 %
